# Primary lung fibroblasts respond to IL-33, IL-13, and IL-17A by secreting factors that activate macrophages

**DOI:** 10.1101/2023.02.28.530495

**Authors:** Jarrett Venezia, Naina Gour, Jeffrey Loube, Wayne Mitzner, Alan L. Scott

## Abstract

There is mounting evidence that macrophage-fibroblast communication is key to the understanding of disease processes. To gain insights into these relationships in the context of progressive lung damage, we measured changes in protein and RNA expression of pulmonary macrophages and fibroblasts upon exposure to IL-33, IL-13, and IL-17A, which are three cytokines often implicated in pathways driving chronic lung remodeling and severe disease like emphysema. Applying an *in vitro* culture system, bulk-RNA sequencing, and protein assays, it was determined that IL-33, IL-13, and IL-17A used alone or in combination activated mouse alveolar macrophages to a modest extent with IL-13 inducing the most vigorous response. While lung fibroblasts also responded modestly to single and paired treatments with IL-33, IL-13, and IL-17A, simultaneous exposure to all three cytokines induced significant activation that was characterized by expression of genes associated with immune cell trafficking and activation, tissue remodeling, and maintenance of the extracellular matrix. Importantly, factors secreted by triple-treated lung fibroblasts resulted in the activation of macrophages *in vitro*. In addition to being the first report describing the cooperative interactions of IL-33, IL-13, and IL-17A on lung fibroblasts, these findings provide additional evidence that fibroblast-macrophage communication is a key component to repair and remodeling in the lung, as well as mechanisms that drive progression of emphysema.

## Introduction

Chronic Lung Disease (CLD) is an umbrella term for disorders that are often the consequence of acute or repeat exposure to environmental toxins, pathogens, or allergens. Well-characterized CLDs include asthma, chronic obstructive pulmonary disease (COPD), and idiopathic pulmonary fibrosis. CLDs pose significant threats to public health; and COPD, specifically, is the third leading cause of death worldwide (WHO, 2022).

CLDs have unique pathologies and prognoses, but considerable overlap exists in terms of the cellular and molecular events thought to drive pathology of each subtype. For example, it has been proposed that macrophages contribute significantly to the damage of lung tissue in asthma, COPD, and pulmonary fibrosis (Bedoret et al., 2009; O’Beirne et al., 2020; Reyfman et al., 2018). Stromal cells have also entered the spotlight, as fibroblast dysregulation has been shown to contribute to the development of asthma (Ingram et al., 2011; Mostaço-Guidolin et al., 2019), COPD (Hallgren et al., 2010; Togo et al., 2008) and pulmonary fibrosis (Kulkarni et al., 2016; Reyfman et al., 2018).

From a molecular standpoint, the cytokines IL-33, IL-13, and IL-17A have all been characterized for their potential association with various CLDs. All three cytokines have been studied extensively for contributing to asthma pathogenesis (Aneas et al., 2021; Grunig et al., 1998; Lajoie et al., 2010; Préfontaine et al., 2009; Wills-Karp et al., 1998). Recently, IL-33 and IL- 17A were recognized for their potential association with pulmonary fibrosis (Li et al., 2014; Mi et al., 2011; Zhang et al., 2019). All three cytokines may also play a role in COPD pathogenesis, as our group and others demonstrated that increased IL-33, IL-13, and IL-17A expression is associated with severe, progressive emphysema (Limjunyawong et al., 2015; Xia et al., 2015).

The goal of the work presented here was to explore the interaction of IL-33, IL-13, and IL- 17A with the resident and recruited lung cells thought to contribute to the progressive lung destruction observed in the elastase-induced experimental emphysema model. Employing transcriptional readouts and protein-based assays, it was determined that primary lung fibroblasts, and not alveolar macrophages, were the cells most responsive to these cytokines and that the most robust response occurred when the fibroblasts were treated with IL-33, IL-13, and IL-17A concurrently. Transcriptional analysis of primary lung fibroblasts showed that triple-treated cells significantly upregulated genes associated with inflammation and tissue- remodeling. These findings also suggested that IL-33, IL-13, and IL-17A induce fibroblasts to secrete factors that influence the activation status of macrophages. Overall, these *in vitro* studies provide the groundwork for further investigating IL-33, IL-13, and IL-17A dynamics and their impact on cellular responsiveness *in vivo*, specifically in the context of chronic lung disease.

## Results

### Alveolar macrophage responsiveness to rIL-13, rIL-33, and rIL-17A *in vitro*

Lung macrophages have been proposed as drivers of emphysema pathogenesis (Hautamaki et al., 1997; O’Beirne et al., 2020; Shibata et al., 2018). Based on observations made in the elastase-induced experimental emphysema (EIEE) model (Limjunyawong et al., 2015) and findings related to other CLDs, like asthma and fibrosis (Aneas et al., 2021; Grunig et al., 1998; Lajoie et al., 2010; Li et al., 2014; Mi et al., 2011; Wijsenbeek et al., 2018; Wills-Karp et al., 1998; Xia et al., 2015; Zhang et al., 2019), we sought to determine whether combinations of IL-33, IL- 13 and IL-17A influence the activation status of lung macrophages. It was postulated that direct exposure of lung macrophages to all three cytokines would cause them to adopt a phenotype capable of driving tissue damage associated with emphysema.

Alveolar macrophages were extracted by bronchoalveolar lavage and stimulated with various combinations of murine rIL-33, rIL-13, and rIL-17A *in vitro* (**Supplemental Table 1**). After 20 hours of cytokine stimulation, test and control alveolar macrophages were processed for expression analysis by RNAseq. Principal component analysis (PCA) of both the raw and normalized data suggested that transcriptional readouts for the rIL-33-only, rIL-17A-only, and rIL- 33/rIL-17A-treated cells overlapped significantly with that of untreated controls (**Figure 1A**). Groups treated with rIL-13, rIL-13/rIL-17A, rIL-33/rIL-13, and rIL-33/rIL-13/rIL-17A all diverged from untreated alveolar macrophages by PCA, suggesting they were activated by cytokine treatment. The groups also overlapped with each other, indicating that cells receiving treatments containing rIL-13 were transcriptionally similar. Differential gene expression (DGE) analysis with DESeq2 was employed for defining unique or overlapping aspects of each treatment group. Relative to baseline, alveolar macrophages receiving only rIL-13 had significant regulation of 972 genes (**Figure 1B**). Several genes associated with alveolar macrophage biology - *Pparg, Ear2, Siglecf, Spp1* (Aegerter et al., 2022; Bain and MacDonald, 2022) - were regulated by rIL-13 in the assay. We also noted differential expression of *Chil3/Ym1, Cd36, Mrc1, Socs1, Cish, Ccl6, Cxcl3,* and *Irf4,* a readout supported by related work characterizing IL4rα-mediated effects *in vitro* (Das et al., 2018; Gordon, 2003; Raes et al., 2005; Scotton et al., 2005). Recombinant IL-13 treatment also regulated genes for tissue remodeling in alveolar macrophages (*Mmp13, Mmp19*) and expression of the IL-33 receptor *St2,* as previously reported (Dagher et al., 2020; Das et al., 2018; Li et al., 2014; Zhang et al., 2021). Consistent with the PCA, the output of the DGE analysis was not statistically significant for rIL-33-only, rIL-17A-only, or rIL-33/rIL-17A-treated cells relative to baseline (**Supplemental Figure 1A**). The addition of rIL-33 had only modest effects on transcription by alveolar macrophages *in vitro* and very few genes were significantly and differentially expressed when the rIL-13-only cells were compared to groups receiving rIL-33 as part of their cocktail (**Supplemental Figure 1B**). Recombinant rIL-17A also had no demonstrable effect on alveolar macrophage transcription *in vitro* (**Supplemental Figure 1A, 1B**) despite detection of *Il17ra* and I*l17rc* transcription in most alveolar macrophage treatment groups (**Figure 1C**). Importantly, there was no indication that IL-33, IL-13 and IL-17A work cooperatively to activate alveolar macrophages (**Supplemental Figure 1C**).

**Figure 1.**
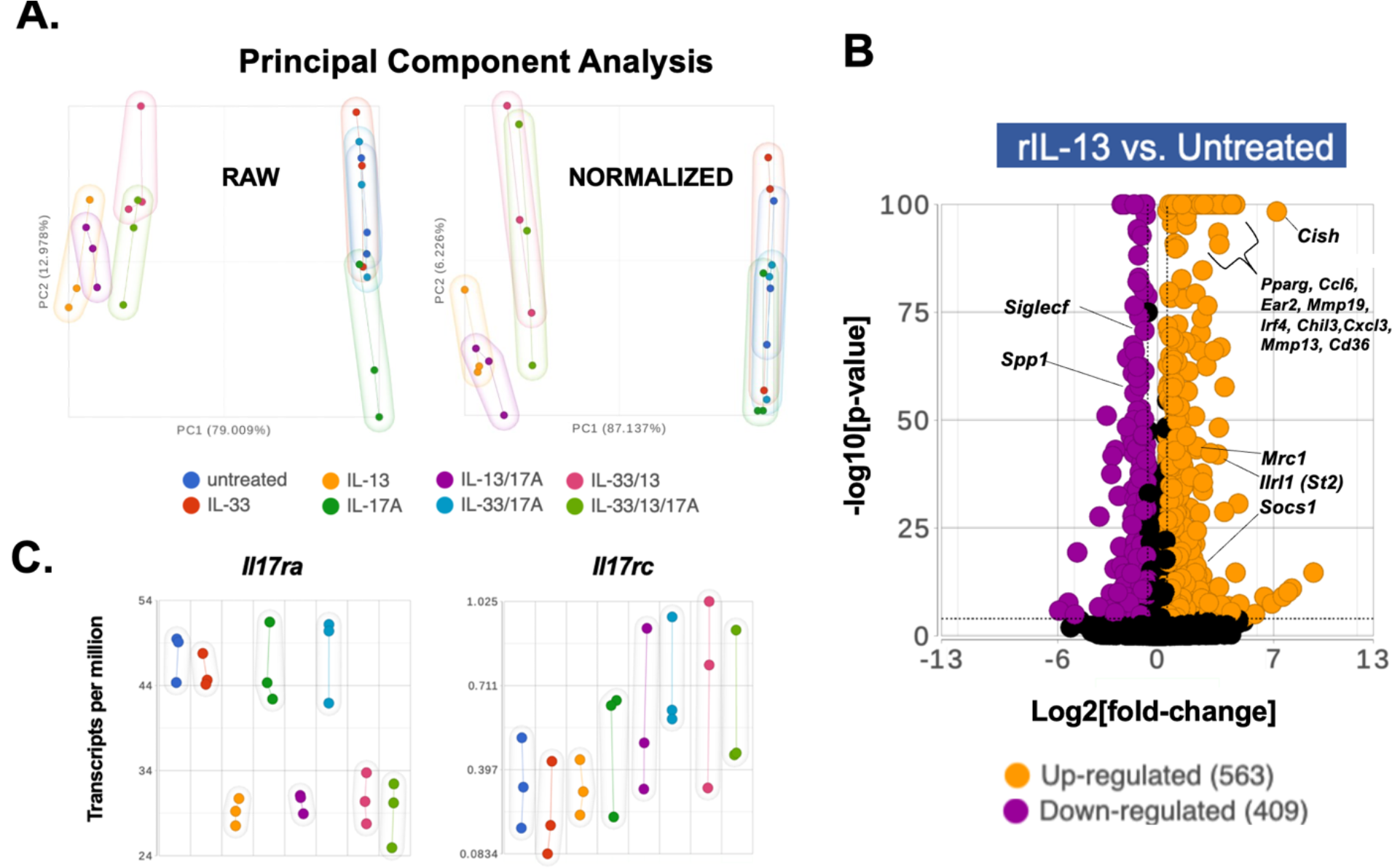
Transcriptomic analysis of alveolar macrophages treated with combinations of recombinant IL-33, IL-13, and IL-17A *in vitro*. Macrophages were extracted from wild-type (WT) BALB/cJ mice as described in **Methods** and were plated in technical triplicate for whole transcriptomic sequencing. (**A**) Principal Component Analysis was used for assessing global, transcriptional changes induced by combinations of recombinant (r) IL-33, rIL-13, and rIL-17A in alveolar macrophages. (**B**) Volcano plot of differentially expressed genes between rIL-13-treated and control alveolar macrophages. Significance cut-offs for this analysis was designated to be p-value <.0001, fold change = +/- 1.5. Gene identifiers were manually annotated and therefore approximated. For volcano plots, the X-axis was transformed as a function of Log2[fold-change], whereas the Y-axis was generated by calculating -log10[p-value]. (**C**) Scatter plots designating expression levels of *Il17ra* and *Il17rc* in alveolar macrophages at baseline and post-cytokine stimulation (TPM-normalized). Bulk RNA sequencing of alveolar macrophages stimulated *in vitro* was performed only once.

Since these *in vitro* results did not support the hypothesis that IL-33, IL-13, and IL-17A work together to directly induce a pathogenic macrophage phenotype, we next explored the alternative hypothesis that IL-33, IL-13, and IL-17A work *<u>indirectly</u>* on alveolar macrophages through the activation of another cell – primary lung fibroblasts. This mechanism is supported by recent evidence that macrophage phenotypes are profoundly influenced by soluble and cognate signals secreted by stromal cells (Buechler et al., 2021; Hou et al., 2019).

### Primary lung fibroblasts responsiveness to rIL-13, rIL-33, and rIL-17A *in vitro*

Primary lung fibroblasts were isolated and established in culture as outlined in the **Methods** and **Figure 2A**. Flow cytometric analysis established that unstimulated primary lung fibroblasts had receptors for both IL-17A and IL-33 deployed on the surface (**Supplemental Figure 2**); and prior reports had established that lung fibroblasts express IL4rα and IL13rα1 (Doucet et al., 1998; Ingram et al., 2004; Saitoa et al., 2003). Lung fibroblasts were incubated with rIL-33, rIL-13, and rIL-17A singly or in combinations (**Supplemental Table 1**) and the culture supernatants were tested for GM-CSF, a growth factor secreted by this cell type (Hamilton, 2019), and one essential for alveolar macrophage growth, development, and function (Gschwend et al., 2021; Guilliams et al., 2013). Combinations that included, at minimum, rIL-33, appeared to elicit modest GM-CSF secretion by fibroblasts *in vitro*; but cells that received all three cytokines produced significantly more GM-CSF than other treatment groups (**Figure 2B**). Reports have suggested that IL-33 regulates GM-CSF secretion by several other cell types (Castro-Dopico et al., 2020; Eissmann et al., 2019; Montanari et al., 2016). To explore a role for IL-33 in activating the lung fibroblasts we had derived *in vitro*, we stimulated cells from *St2*^-/-^ mice, which cannot mediate IL-33 signaling, and compared their GM-CSF response to that of WT fibroblasts upon exposure to rIL-33/rIL-13/rIL-17A. Unlike WT cells, *St2*^-/-^ fibroblasts did not produce significant levels of GM-CSF upon triple-treatment (**Figure 2C**). Taken together, these preliminary findings suggested that IL-33 was the dominant signal driving GM-CSF production *in vitro*, but IL-33, IL-13, and IL-17A may cooperatively regulate GM-CSF production in primary lung fibroblasts.

**Figure 2.**
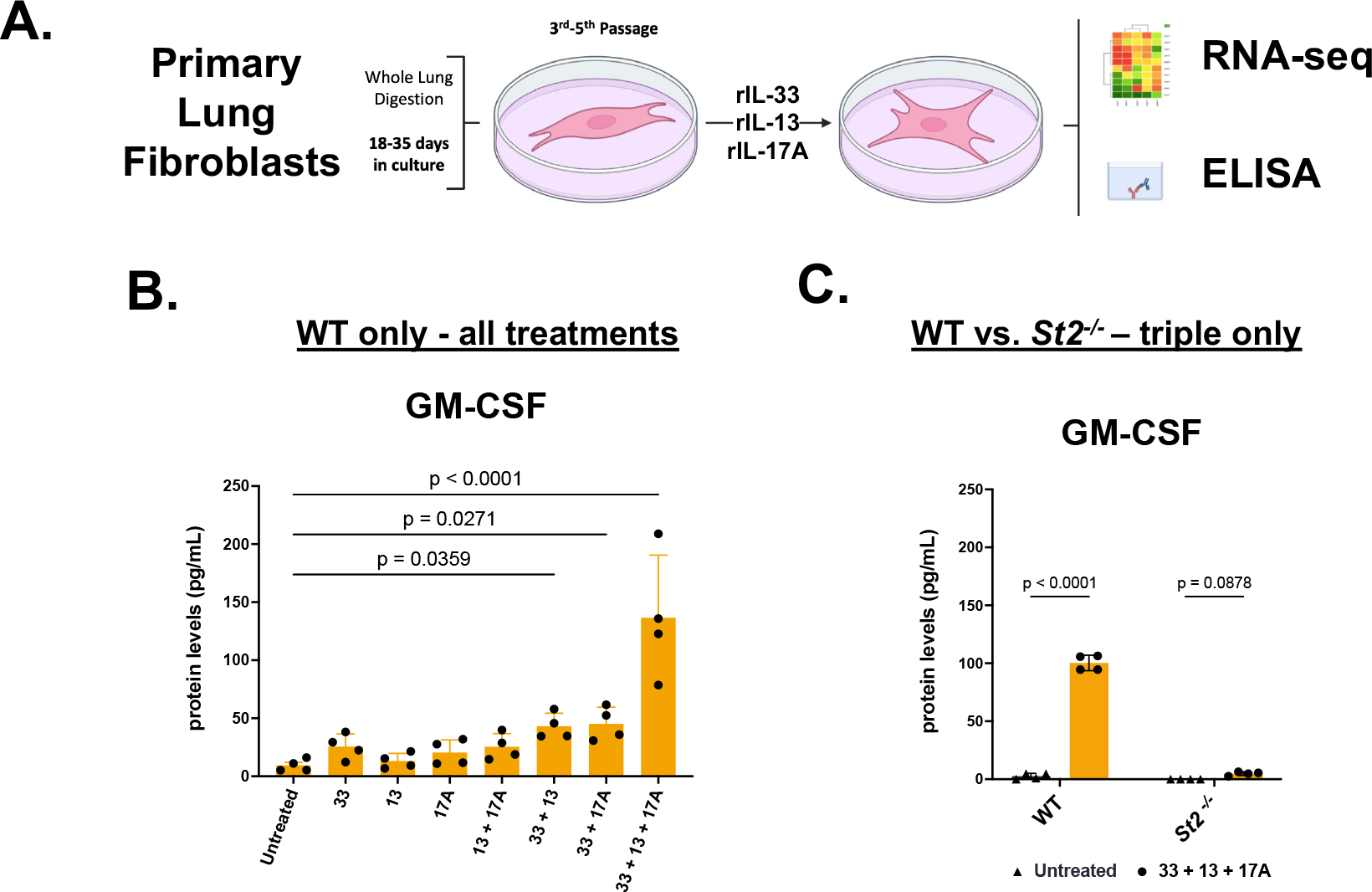
GM-CSF production by primary lung fibroblasts induced by combinations of recombinant IL-33, IL-13, and IL-17A (**A**) Lung fibroblasts first were passaged 3 to 5 times and then plated in quadruplicate for preliminary stimulation assays, as described in **Methods**. (**B**) Levels of GM-CSF (pg/ml) secreted into the culture supernatant by wild-type (WT) primary lung fibroblasts treated with combinations of recombinant (r) IL-33, rIL-13, and rIL-17A. (**C**) Levels of GM-CSF (pg/ml) secreted into the culture supernatant by WT and *St2^-/-^* fibroblasts treated with rIL-33/rIL-13a/rIL-17A *in vitro*. Significance (<u>p-value</u> < <u>.05</u>) was determined with ordinary one-way (**B**) or two-way ANOVA (**C**). Multiple comparison testing was computed post-hoc, and p-values were calculated without mathematical correction using the unprotected Fischer’s Least Significant Difference test. (**B**) <u>rIL-</u> <u>33/rIL-13/rIL-17A vs. untreated, p <0.0001</u>; <u>rIL-33/rIL-17A vs. untreated, p =0.0271</u>; <u>rIL-33/rIL-13,</u> <u>p =0.0359</u>. (**C**) <u>WT, p <0.0001</u>; *St2^-/-^*, p =0.0878. Experiments for (**B**) and (**C**) were performed at least twice, yielding similar outcomes.

To define additional gene products induced by rIL-33, rIL-13, and rIL-17A *in vitro*, we performed bulk RNAseq on WT lung fibroblasts exposed to cytokine treatments that contained, at minimum, rIL-33 (**Supplemental Table 2**) and compared expression profiles to untreated cells. Global transcriptional differences within and between groups were computed in an unsupervised manner with principal component analysis (PCA) and hierarchical clustering (hclust). In contrast to what was observed for alveolar macrophages (**Figure 1A**), all treatment groups clustered distinctly from each other and from the untreated controls in PCA space (**Figure 3A**). These patterns offered preliminary support for the idea that combinations of rIL-33/rIL-13/IL-17A induce unique activation phenotypes in primary lung fibroblasts. Illustrated by heatmap, hclust suggested that untreated and rIL-33-treated fibroblasts had similar expression profiles (**Supplementary Figure 3**); but the remaining treatment groups (rIL-33/rIL-13, rIL-33/rIL-17A, and rIL-33/rIL-13/rIL-17A) had global signatures that were clearly different from baseline and/or rIL- 33-treated cells. To dissect subtle transcriptional changes and common features that exist in double- and triple-treated fibroblasts, we performed differential gene expression (DGE) analysis by comparing each treatment group to untreated cells with DESeq2. DGE analysis provided support for the idea that IL-13, IL-33, and IL-17A signaling work cooperatively to enhance production of GM-CSF by primary lung fibroblasts, as transcriptional regulation of *Csf2* (**Figure 3B**) correlated with the observations made at the protein level (**Figure 2B**). DGE analysis of rIL- 33/rIL-13/rIL-17A-treated fibroblasts versus untreated control cells revealed 836 significantly upregulated and 806 significantly downregulated genes (1,642 genes total; **Figure 3C**). Genes differentially expressed include those encoding inflammatory and tissue-remodeling proteins like *Il33, Tslp, Cxcl2, Mmp10, Mcam, Acta2, Col4a2.* When compared to untreated cells, fibroblasts treated with rIL-33/rIL-13 or rIL-33/rIL-17A had significant differential expression in 1,536 and 1,228 genes, respectively (**Figure 3D, 3E**); while treatment with rIL-33 had induced differential expression in 473 genes total (**Figure 3F**).

**Figure 3.**
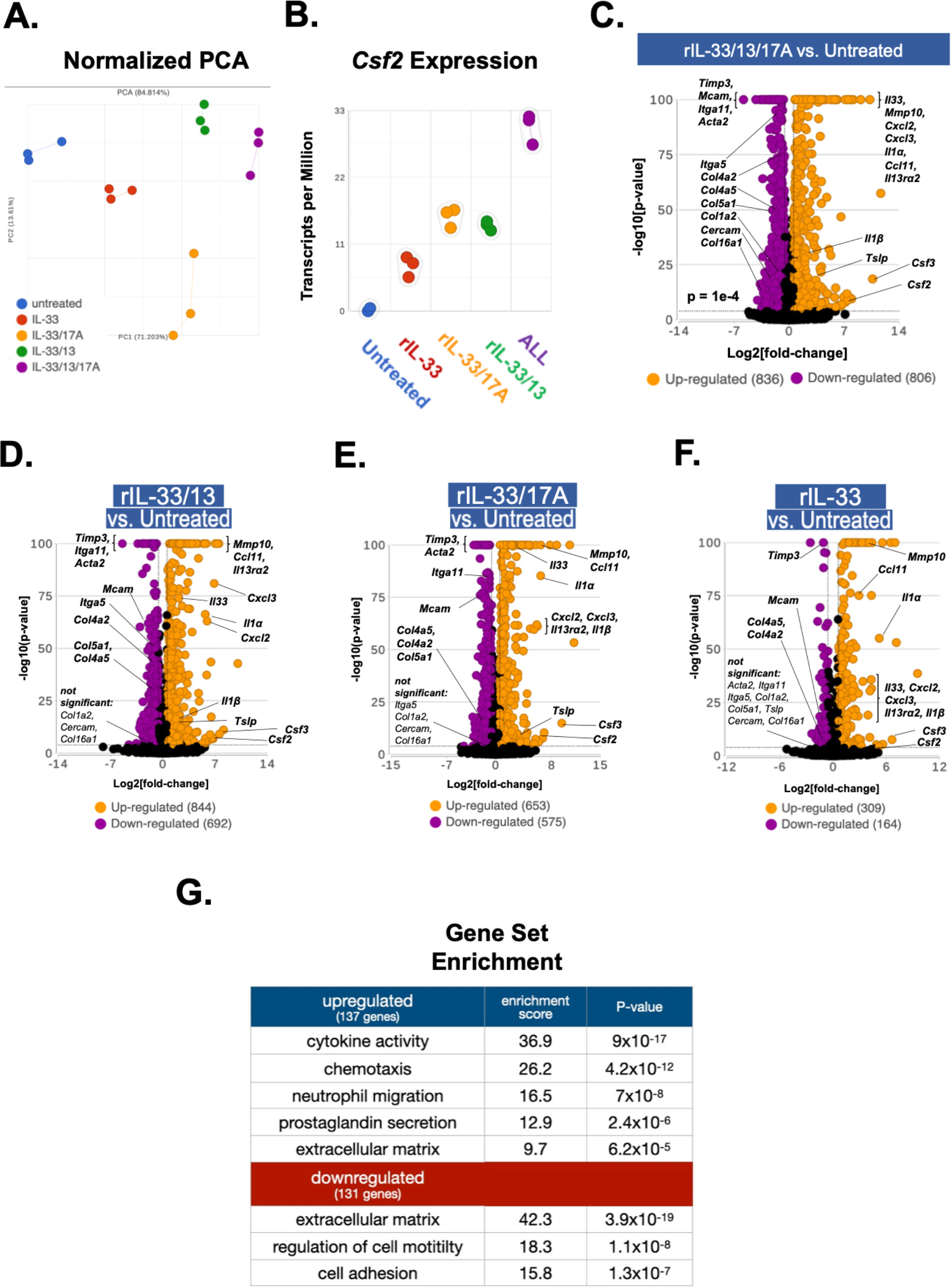
Whole transcriptomic analysis of primary lung fibroblasts treated with combinations of recombinant IL-33, IL-13, and IL-17A *in vitro*. Fibroblasts were cultured and stimulated in triplicate with combinations of cytokines, as described in the **Methods**. (**A**) Principal component analysis of the transcriptional changes induced by four cytokine combinations *in vitro*. (**B**) *Csf2* expression in primary lung fibroblasts determined by RNAseq and illustrated by scatter plot (TPM-normalized). (**C-F**) Volcano plots showing differential gene expression of cells treated with recombinant (r) cytokines versus untreated fibroblasts; (**C**) <u>rIL-33/rIL-13/rIL-17A,</u> (**D**) <u>rIL-33/13</u>, (**E**) <u>rIL-33/17A</u>, (**F**) <u>rIL-33 vs.</u> <u>untreated</u>. Significance cut-offs for this analysis was designated to be p-value <.0001, fold change +/- 1.5. Gene identifiers in each volcano plot were manually annotated and therefore approximated. For generating volcano plots, data were transformed as function of -log10[p-value] (Y-axis) and Log2[fold-change] (X-axis). (**G**) Gene Set Enrichment (GO analysis) of the top 238 differentially expressed genes identified in triple-treated cells. Bulk RNA sequencing of primary lung fibroblasts stimulated *in vitro* was performed only once.

Of the 1,642 statistically significant features (p<.0001) that were differentially expressed by triple-treated fibroblasts relative to untreated cells, only 268 genes remained significantly different when triple-treated fibroblasts were also compared to those stimulated with rIL-33/rIL- 13, rIL-33/rIL-17A, or rIL-33 alone (features were eliminated if the comparison yielded a false discovery rate > .01; **Supplemental Figure 4**). Gene ontology (GO) analysis (Ashburner et al., 2000) was performed on this gene set (**Figure 3G**), revealing significant fold enrichment for upregulated genes associated with cytokine activity (*Il1a, Il6, Tslp, Il33, Il1b, Il11*) and neutrophil trafficking (*Cxcl1, Cxcl2, Cxcl3, Cxcl5*), prostaglandin production (*Ptges, Ptgs2*), and enzymes that modify the extracellular matrix (*Mmp10, Mmp13, Mmp12*). The GO analysis also identified that genes associated with the synthesis of extracellular matrix (*Col6a3, Col11a1, Col4a5, Col15a1, Tgfb3, Lama2, Vcl*) and the regulation of cell motility and adhesion (*Itga11, Itga5, Mcam, Pcdh19, Fbln5*) were downregulated (**Figure 3G, Supplemental Figure 4**). Differentially expressed genes from this list were grouped into functional categories to illustrate how triple-treatment impacted fibroblast activation when compared to cells stimulated with rIL-33, rIL-33/rIL-17A, or rIL-33/rIL- 13 (**Figure 4**). Altogether, the triple-treated fibroblasts significantly upregulated genes for immune cell activation and tissue remodeling, but downregulated genes for extracellular matrix synthesis (**Figure 4**, **Supplemental Figure 4**).

**Figure 4.**
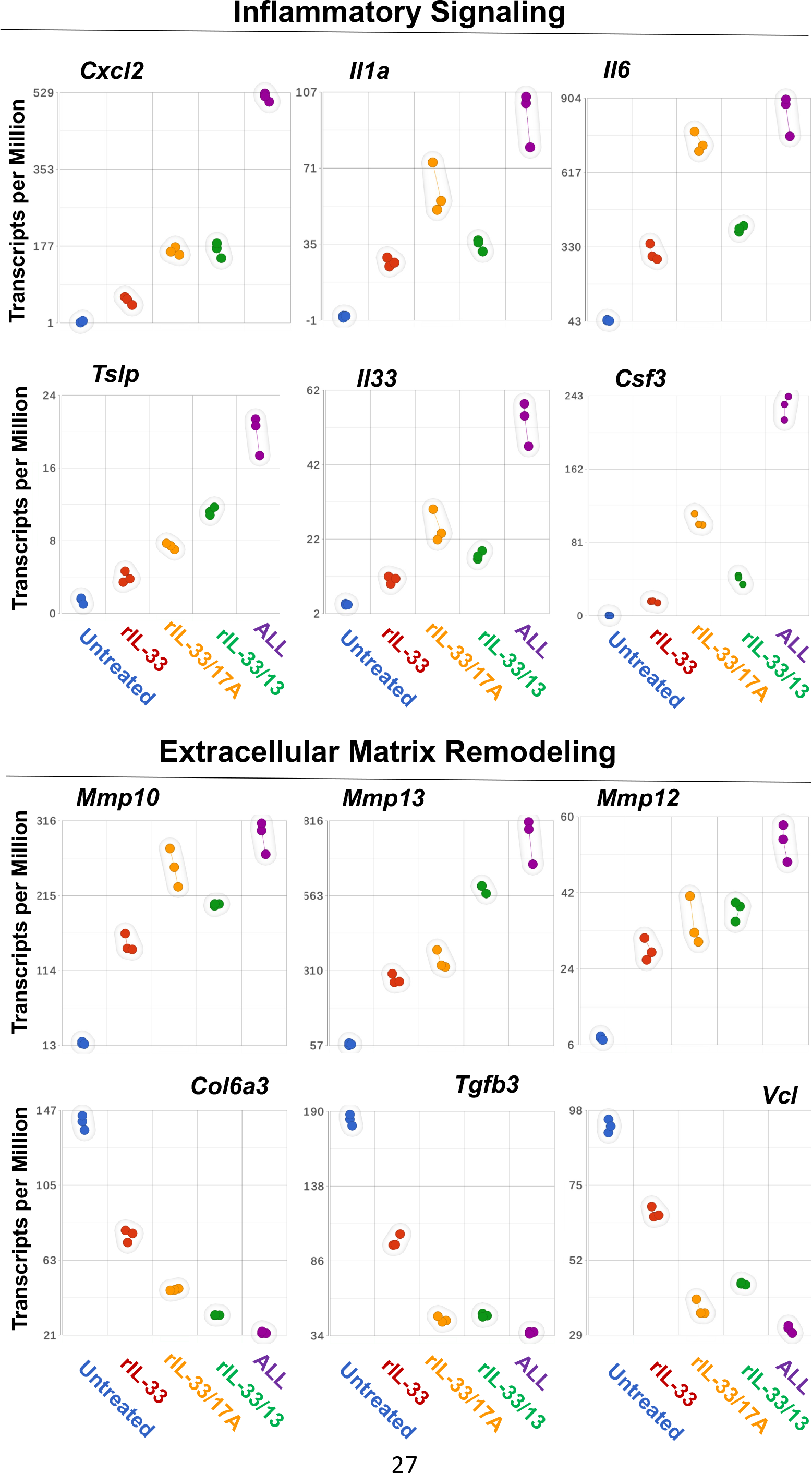
Direct comparison of primary lung fibroblast absolute transcript abundance at baseline and post-cytokine stimulation. Genes that were differentially and significantly expressed by triple-treated fibroblasts were subsampled for representation by 2D scatter plot. Each treatment group is portrayed on the X-axis; the Y axis represents transcript abundance (TPM-normalized) for the following: *Cxcl2, Il1a, Il6, Tslp Il33, Csf3, Mmp10, Mmp13, Mmp12, Col6a3, Tgfb3, Vcl*. These genes are among the top 268 features that were determined by independently comparing triple-treated fibroblasts to each group in the assay. Fold-change values for these comparisons, which were determined to be significant for having a false-discovery rate of < 0.01, are available for reference in **Supplemental Table 4**.

### Functional Implications for primary lung fibroblast stimulation by rIL-33, rIL-13, and rIL-17A

To test the idea that rIL-33/rIL-13/rIL-17A stimulation induced lung fibroblasts to secrete factors that have the capacity to activate macrophages, we employed an *in vitro* assay for stimulating bone marrow-derived macrophages (BMDM) with fibroblast-conditioned media (“FCM”; **Figure 5A**). Independently cultured, primary lung fibroblasts were stimulated with combinations of rIL-33, rIL-13, and rIL-17A for 20 hours. Fibroblasts were then rinsed several times to wash away recombinant proteins, replenished with fresh media, and cultured for an additional 20 hours to generated conditioned medium. Sterile-filtered FCM was subsequently transferred to BMDM in culture medium at 20% v/v, and macrophages were incubated for 24 hours before harvesting total RNA for RT-qPCR analysis. The genes assayed by RT-qPCR (*Arg1*, *Ym1, Retnla*) were chosen for representing M2 activation, which is the macrophage phenotype associated with wound healing and tissue-remodeling (Ley, 2017; Mills et al., 2000). FCM derived from rIL-33/rIL-17A and rIL-33/rIL-13-treated fibroblasts each induced a significant increase in *Arg1* expression by BMDM (**Figure 5B**). However, these elevations were relatively low compared to the robust *Arg1* induction in macrophages that received FCM from triple-treated fibroblasts. These data were significant in the context of a more conventional approach to macrophage activation *in vitro*, during which BMDM were directly treated with rIL-13, as opposed to FCM, and compared to cells treated with recombinant IL-33/IL-13/IL-17A (**Figure 5C**). In these assays, triple-treatment regulated *Arg1* expression in a manner comparable to BMDM treated with rIL-13 alone – which, along with IL-4, is a known and potent inducer of *Arg1* in cultured macrophages (Gordon, 2003; Ley, 2017; Mills et al., 2000; Raes et al., 2005). Interestingly, BMDM treated with FCM activated *Arg1* in the absence of other M2 genes like *Ym1* and *Retnla* (**Supplemental Figure 5A**, **5B**). As expected, though, the BMDM derived *in vitro* expressed *Retnla* in response to direct stimulation with rIL-13 and/or triple-treatment as part of the M2 phenotype (**Supplemental Figure 5C**). These findings are supportive of the notion that lung fibroblasts stimulated with rIL- 13, rIL-33, and rIL-17A secrete molecules with potential to influence the activation status of macrophages.

**Figure 5.**
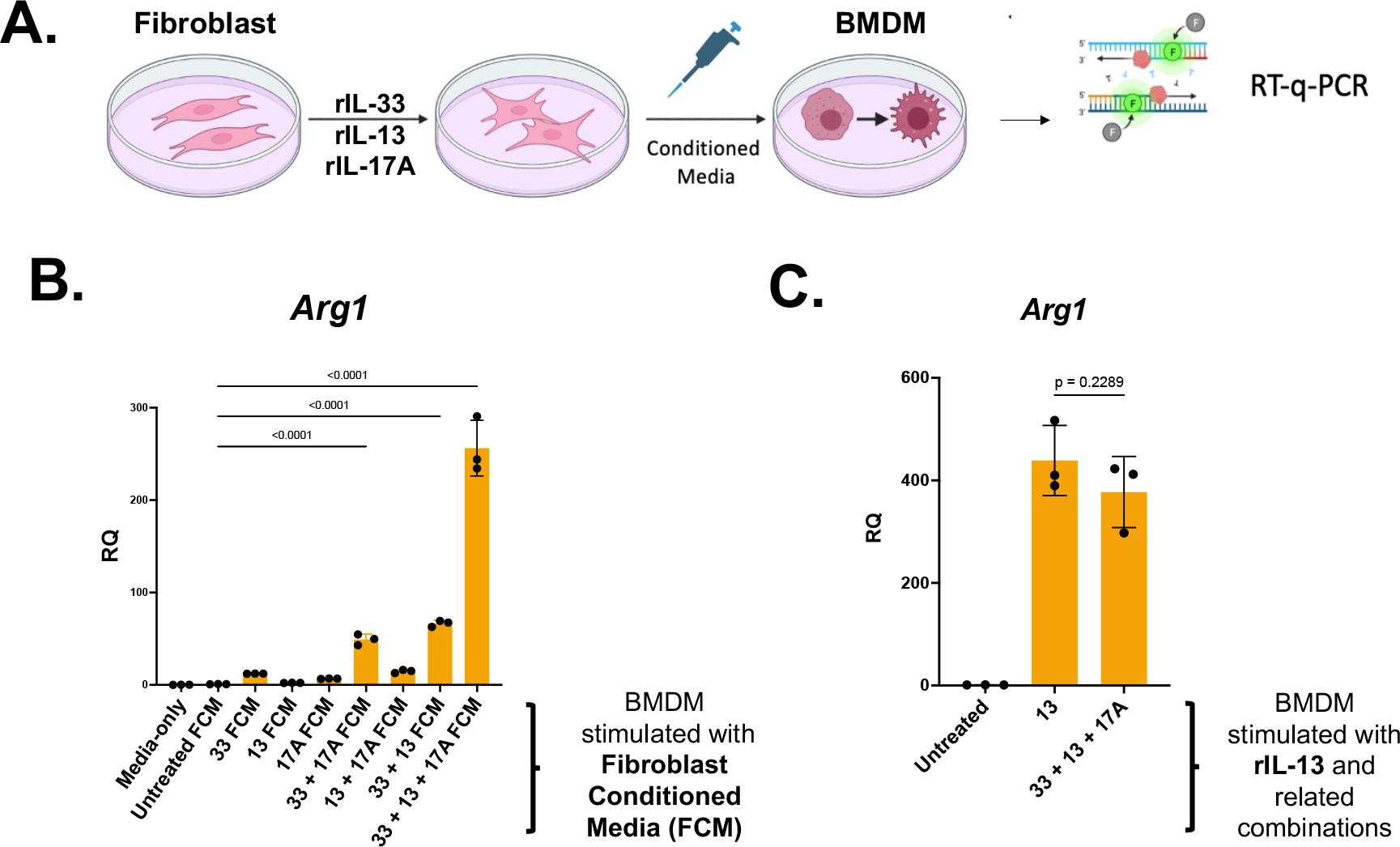
Fibroblast conditioned medium activates bone-marrow derived macrophages *in vitro*. Fibroblasts were isolated from digested whole lung, whereas macrophages (BMDM) were derived *in vitro* from monocytes extracted from the bone marrow. (**A**) Independent cultures of fibroblasts were subjected to cytokine treatment, but for the purpose of harvesting fibroblast- conditioned media (FCM) that can be transferred to BMDM cultures. Activation phenotypes for BMDM were assessed by RT-qPCR. (**B**) Induction of *Arg1* expression in BMDM cultures activated by FCM. (**C**) Induction of *Arg1* expression in BMDM cultures treated with cytokine cocktails containing rIL-13. (**B-C**) Significance (<u>p-value</u> < <u>.05</u>) was determined with ordinary one-way ANOVA. Multiple comparison testing was computed post-hoc, and p-values were calculated without mathematical correction using the unprotected Fischer’s Least Significant Difference test. (**B**) <u>rIL-33/rIL-13/rIL-17A FCM vs. untreated FCM, p <0.0001</u>; <u>rIL-33/rIL-13 FCM vs. untreated</u> <u>FCM, p <0.0001</u>; <u>rIL-33/rIL-17A FCM vs. untreated FCM, p <0.0001</u>. (**C**) rIL-13 vs. rIL-33/rIL-13/rIL-17A, p =0.2289. (**B-C**) Data representative of an experiment performed at least twice, yielding comparable outcomes.

## Discussion

Previous work indicated that IL-33, IL-13 and IL-17A might work together to induce lung macrophages to take on a phenotype that mediates progressive tissue damage (Limjunyawong et al., 2015). In the context of elastase-induced experimental emphysema (EIEE), a rapid increase in IL-33 occurs post-initiation of disease followed by upregulated expression of IL-13 and IL-17A. The concomitant increases in pulmonary IL-33, IL-13, and IL-17A were temporally associated with lung macrophage activation and the onset of progressive degradation of alveoli in this experimental model. Several reports already pointed to a role for these cytokines in macrophage activation in the context of inflammation (Barin et al., 2012; Dagher et al., 2020; Ge et al., 2014; Li et al., 2014). Previous work demonstrated that IL-33 synergizes with IL-13 for modulating macrophage effector functions *in vitro* and *in vivo* during wound repair (Dagher et al., 2020; He et al., 2017; Kurowska-Stolarska et al., 2009), but to our knowledge there are no reports on the combined effects of IL-33/IL-13/IL-17A treatment on immune or stromal cells.

Under the conditions used for this study, treatment of alveolar macrophages with rIL-13 induced significant changes in gene expression while the addition of rIL-33 or rIL-17A had little to no effect on transcriptional signatures (**Figure 1**) and did not induce an expression profile consistent with a tissue-destructive phenotype. Since these results did not support a model for direct stimulation, we were prompted to consider other mechanisms by which IL-33, IL-13 and/or IL-17A activate alveolar macrophages. In two chronic-progressive disease models in which tissue-resident macrophages are implicated in the mechanism of pathogenesis – pulmonary fibrosis and myocarditis – fibroblasts have been identified as key accessory cells that contribute to the phenotypic changes in macrophages (Hou et al., 2019; Reyfman et al., 2018). Thus, we tested the hypothesis that lung fibroblasts are a primary target of IL-33, IL-13, and IL-17A and that stromal cells produce molecules that function in the reprogramming of alveolar macrophages into pathogenic cells.

An important finding from these studies was that rIL-33, rIL-13, and rIL-17A appear to work in a cooperative fashion to activate primary lung fibroblasts at the protein (**Figure 2**) and transcriptional levels (**Figure 3**), but enhanced GM-CSF transcription and secretion was just one facet of a more complex activation phenotype. Bulk RNAseq revealed significant, differential gene expression for inflammatory signaling, tissue remodeling, and extracellular matrix proteins by triple-treated fibroblasts (**Figure 4**). GM-CSF in the lungs is necessary for alveolar macrophage development *in vivo* (Gschwend et al., 2021; Guilliams et al., 2013); and lack of *Csf2* expression or autoimmune responses that block GM-CSF function results in dysregulated alveolar macrophages and diseases such as pulmonary alveolar proteinosis (Uchida et al., 2007). Although all three cytokines were required for maximal production of GM-CSF in our *in vitro* system, lack of GM-CSF secretion by *St2^-/-^* lung fibroblasts clearly indicated that IL-33-mediated signaling was a major contributor to the phenotype.

Compared to baseline expression, triple-treated fibroblasts downregulated genes typically expressed by myofibroblasts (**Figure 4, Supplemental Figure 4**), which are activated stromal cells native to several tissues, including the lung. Myofibroblasts robustly express *Acta2* as well as genes for extracellular matrix synthesis (Foster et al., 1990; Kulkarni et al., 2016; Tomasek et al., 2002); and primary lung fibroblasts generated by our *in vitro* protocol are known to develop myofibroblast-like characteristics at baseline with time and passage (Edelman and Redente, 2018; Foster et al., 1990; Seluanov et al., 2010). Upon triple-treatment, however, these fibroblasts appear to significantly downregulate *Acta2* and key genes *(i.e., Col6a3, Col11a1, Col4a5, Col15a1, Lama2, Fbln5*) that code for proteins highly abundant in the murine lung extracellular matrix (Burgstaller et al., 2017).

Recently, it was suggested that nuclear IL-33 may restrain expression of several fibrotic gene sets in dermal fibroblasts (Gatti et al., 2021). On the other hand, pro-fibrotic effects have been reported in human or murine lung fibroblasts stimulated with IL-13 or IL-4 alone *in vitro* (Doucet et al., 1998; Firszt et al., 2014; Hashimoto et al., 2001; Ingram et al., 2004; Saitoa et al., 2003). Outside of the scope of pulmonary fibrosis (Mi et al., 2011; Zhang et al., 2019), IL-17A signaling in lung fibroblasts has not been extensively studied. In our work, triple-treatment elicited an intriguing transcriptional phenotype *in vitro* that may begin to explain how lung fibroblasts contribute to lung destruction and emphysema pathogenesis, which is characterized by progressive degradation of alveoli and a lung microenvironment that fails to regenerate (Foster et al., 1990; Kulkarni et al., 2016).

Finally, several genes upregulated in triple-treated fibroblasts encode products with the potential to mediate activation of macrophages. To broadly characterize this putative activation circuit, we demonstrated that fibroblast conditioned media from triple-treated lung fibroblasts induced macrophage expression of *Arg1.* Conventionally, expression of *Arg1* is strongly associated with IL-4- and/or IL-13-mediated activation of M2 macrophages (Das et al., 2018; Gordon, 2003; Raes et al., 2005); but expression is now thought to be inducible by other factors, including GM-CSF (Su et al., 2021). As the only protein confirmed to be secreted by fibroblasts with ELISA (**Figure 2**), GM-CSF signaling may begin to explain BMDM upregulation of *Arg1* in response to FCM *in vitro*. In support of this theory, we indeed confirmed that BMDM generated for these assays deploy at least one component of the GM-CSF receptor on their surface prior to stimulation with FCM (**Supplemental Figure 5D**). Unlike rIL-4/rIL-13 stimulation, FCM might regulate expression of *Arg1* without affecting the transcription of other, common M2 genes, such as *Retnla* and *Ym1.* While additional work is necessary for determining the nature and scope of lung fibroblast-induced macrophage activation *in vivo*, these *in vitro* observations provide a foundation for investigating how stromal cells communicate with macrophages during an injury response in the lungs.

## Materials & Methods

### Mice

All experiments were conducted in accordance with the standards established by the United States Animal Welfare Acts, set forth in NIH guidelines and the Policy and Procedures Manual of the Johns Hopkins University Animal Care and Use Committee. Animals were maintained in filter-topped cages at 72°C, 50-60% humidity with 14:10-hour light/dark cycle and *ad libitum* access to food and water. Cells for *in vitro* studies were derived from mice on the BALB/c background. Cells were harvested from either males or females and pooled for *in vitro* culture. Wild-type (WT) BALB/cJ were originally purchased from Jackson Laboratories (#000651). Mice were bred and housed in the Johns Hopkins School of Public Health animal facilities. *St2^-/-^* mice (Andrew N.J. McKenzie, Cambridge, U.K.) were also bred in-house for these studies

### Recombinant Cytokines

Recombinant (r) mouse cytokines were purchased from BioLegend: rIL-33 (580502), rIL- 13 (575904), rIL-17A (576002). The optimal concentration of each recombinant cytokine was determined in preliminary studies. The same concentrations were used for all *in vitro* stimulations: 30 ng/mL (rIL-33), 20 ng/mL (rIL-13), 30 ng/mL (rIL-17A).

### Primary lung fibroblast cell cultures

BALB/c mice (3-6 weeks old) were anesthetized by intraperitoneal (IP) injection with a mixture (1:1) of ketamine (100 mg/kg) and xylazine (10 mg/kg) in sterile water. Intact, non-perfused lungs were extracted, finely minced with a razor blade and placed on ice in RPMI 1640

(Corning 10-40-CV). Liberase TL (Roche 5401020001) and DNase I (Sigma DN25) in RPMI, was added to each homogenate, and samples were incubated for 45 minutes at 37°C and 5% CO2. Post-digestion, homogenates were passed through a 40 μm cell strainer and rinsed with RPMI, centrifuged, and suspended in 1% Penicillin/Streptomycin (Gibco, 15140122) + 10% heat-inactivated fetal bovine serum (“FBS”; Corning, 35-010-CV) in Dulbecco’s Modified Eagle’s Medium (“DMEM”; Corning, 10-017-CV). For the expansion and maintenance of primary lung fibroblasts, lung homogenates were directly plated in culture media. With time and at least 3 rounds of passage, lung-derived fibroblasts become the dominant population in the cultures (Edelman and Redente, 2018; Foster et al., 1990; Seluanov et al., 2010). The medium was changed every other day, and cell passage was performed at 80% confluency. Only cells from passage 3-5 were used for experiments.

### Primary Alveolar Macrophage cultures

BALB/cJ mice (6-12 weeks old) were euthanized with ketamine/xylazine, a tracheostomy was performed, and an 18G canula was inserted. Bronchoalveolar lavage (“BAL”) was performed as previously described (Busch et al., 2019) using the BAL buffer (2 mM EDTA + 1% heat inactivated FBS in Dulbecco’s phosphate buffered saline (“dPBS”). 900 μl pre-warmed (37°C) BAL buffer was introduced via the canula and slowly drawn back from the lungs with a recovery of 500 μl to 800 μl. Cells were rinsed with cold BAL buffer and directly plated in culture media (1% Penicillin/Streptomycin + 10% heat-inactivated FBS in DMEM). After 20 minutes in standard culture conditions (37°C, 5% CO2), plates were rinsed with dPBS to remove nonadherent cells and replenished with fresh culture media. Alveolar macrophages were in culture for no longer than 24 hours. The purity and viability of alveolar macrophages were estimated by light microscopy and trypan blue staining.

### mRNA Preparation and cDNA synthesis

Total RNA from fibroblasts or macrophages was prepared using commercially available RNeasy kits and protocols (Qiagen 74004 or 74104). For RT-PCR analysis, total RNA was reverse transcribed into cDNA with LunaScript RT Supermix (New England Biolabs, E3010) according to manufacturer’s protocol. RNA was stored at –80°C prior to use; and cDNA was stored at 4°C.

### RNA Expression Analysis

Alveolar macrophages were isolated, as described above, and a total of 2.25 x 10^5^ cells per well were seeded in 24-well assay plates (Corning 3524) for stimulation with combinations of recombinant cytokines (**Supplemental Table 1**). Primary lung fibroblasts were also isolated, as described above, and 1.2 x 10^6^ cells per well were seeded in 6-well assay plates (Corning 3516) and treated with combinations of recombinant cytokines (**Supplemental Table 2**). For both cell types, total RNA was isolated after 20 hours of stimulation with the recombinant cytokines and the column-purified RNA was sent to Novogene for QA/QC, library preparation, and generation of paired-end reads (150 bp) using the NovaSeq 6000 Sequencing System (Illumina). Computational analysis was performed using Partek Flow (St. Louis, MO) with .bam files that were previously aligned by Novogene using either the HISAT2 or STAR algorithms (Kim et al., 2019; Mortazavi et al., 2008). A *Mus musculus* genome (‘mm10’ a.k.a. GRCm38) was referenced for alignment. Mapped reads were subjected to standard RNAseq QA/QC to assess for nucleotide mismatches or library size differences between samples. Low quality reads were excluded from the dataset for having a Phred II index score of < 30 (Ewing and Green, 1998; Ewing et al., 1998), and the remaining alignments were quantified with the Ensembl release 102 annotation model (Li and Dewey, 2011; Zhao and Zhang, 2015). Count matrices were then processed for data visualization and biological inference downstream. Principal component analysis, hierarchical clustering, and scatter plots were employed for visualizing expression data. Differential gene expression (“DGE”) analysis was executed using Partek Flow with DESeq2 (Anders and Huber, 2010; Love et al., 2014) and visualized with volcano plots. For DGE analysis, raw counts were normalized by the ‘median ratio’ method in DESeq2 by default. For visualizing absolute gene expression by scatter plot, ‘Transcripts per million’ normalization was applied.

### Bone Marrow-Derived Macrophage (BMDM) Cultures

BALB/cJ mice (6-12 weeks old) were euthanized as outlined above. The long bones were removed, their surface washed with 70% ethanol, and rinsed in dPBS. Bone marrow was flushed with DMEM using a syringe fitted with a 28 G needle and the cells were passed through a 40 μm cell strainer. Cells were rinsed with DMEM, centrifuged, and plated directly in bone marrow monocyte differentiation media (1% Penicillin/Streptomycin + 10% Heat inactivated-FBS + 20% L-cell conditioned media in DMEM). Medium was replaced every 3 days and the BMDM were used after 7 days of culture in the differentiation media.

### Generation of Fibroblast Conditioned Media

Primary lung fibroblast cultures were treated with combinations of rIL-33, rIL-13, and rIL- 17A for generating fibroblast conditioned media (“FCM”). Fibroblast-only cultures were stimulated with cytokine combinations for 20 hours, rinsed three times with dPBS to remove residual recombinant cytokines, replenished with media, and incubated (37°C, 5% CO2) for an additional 20 hours as the source of FCM. Supernatants were collected, sterilized with a 0.22 μm filter, and transferred to BMDM cultures (1 x 10^6^ cells per well) at 20% v/v in DMEM (1% Penicillin/Streptomycin + 10% heat-inactivated-FBS). After 24 hours of exposure to FCM, total RNA was collected from BMDM, as described above.

### Enzyme-Linked Immunoassay (ELISA) for measuring secretion of murine GM-CSF

Commercially available kits and manufacturer protocols from R&D Systems (mCsf2; DY416) were employed for ELISA. Unless clearly stated otherwise, 25,000 cells were seeded in 96-well assay plates for cytokine treatment. At the conclusion of each experiment, supernatants were collected and stored in low-binding 96-well plates (Greiner 655904) at –80°C. Protein concentrations were measured from supernatants by ELISA using a fluorescent microplate reader, directly comparing the signal from each sample to that of a standard curve.

### Reverse Transcription – Quantitative Polymerase Chain Reaction (RT-qPCR)

Purified RNA was quantified using a Qubit 2.0 (Thermo Fisher, Q32886) and reverse transcribed, as described above. TaqMan-based qPCR was performed using EagleTaq Universal Master Mix (Roche, 7260288190), and data were acquired on a StepOnePlus (Applied Biosystems, 4376600) with the following fluorescently-labeled probes: *Arg1* (Thermo Fisher*; Mm00475988_m1), Rps14 (Mm00849906_g1), Ctsa (Mm00447197_m1), Retnla (Mm00445109_m1), Ym1 (Mm00657889_mH), Gapdh (Mm99999915_g1).* A commercial algorithm (RQ, Applied Biosystems) was used for determining expression values, or ‘relative quantitation’, of several genes. This computation is similar to the 2^-ΔΔCt^ method (Livak and Schmittgen, 2001). For calculating relative quantitation, expression of *Rps14, Ctsa* and *Gapdh* were acquired by qPCR in parallel as controls (i.e. ‘housekeeping genes’).

### Flow Cytometry

Cultured lung fibroblasts were lifted by incubating cells with Accutase (BioLegend 423201) for up to 20 minutes at room temperature. In preparation for cytometry, fibroblasts were suspended in dPBS (for vitality dye only) or FACS Buffer, consisting of 2 mM EDTA (Corning, 46- 034-CL) + 2% heat-inactivated FBS in dPBS. Cells were treated with Fc block (anti-CD16/CD32; clone 2.4G2, BioXcell) and staining was performed with combinations of the following: LIVE/DEAD Aqua (L34957; eBioscience), anti-mouse IL17ra-APC (clone 657643, R&D systems), anti-mouse T1/St2-FITC (clone DJ8, mdbioproducts), anti-mouse Csf2ra-Alexa 700 (clone 698423, R&D systems), or anti-mouse CD11b-BV785 (clone M1/70, BioLegend). Single-fluorophore controls were prepared using either cytometry beads (Becton Dickinson 552843) or cells isolated on the day of experimentation. Acquisition was performed on an LSR-II (Becton Dickinson) using the FACSDiva software. Compensation was finalized in FlowJo (Becton Dickinson), and this software was also used for data analysis and visualization. For all flow cytometry experiments, unstained and fluorescence-minus-one control samples were employed for analysis and for determining gates.

### Statistical Analysis and Illustrations

For ELISA and RT-q-PCR readouts, one- or two-way ANOVA was performed in GraphPad Prism (San Diego, CA), which was also used for visualization of these data. For multiple-comparison testing, uncorrected p-values were computed using the unprotected Fischer’s Least Significant Difference test, as per statistical recommendations in GraphPad Prism. For RNAseq, all statistics were calculated with computational tasks in Partek Flow (St. Louis, MO). Differential analysis with DESeq2 computed an ANOVA table, per comparison, with the following statistics: fold-change, p-value, false-discovery rate step-up. BioRender was also used for several visuals.

### Data Availability

Alveolar macrophage and primary lung fibroblast RNAseq data will be submitted to the Gene Expression Omnibus (NCBI) upon publication. Raw ELISA, RT-q-PCR, and flow cytometry data are available upon request.

---

**Supplemental Figure 1.**
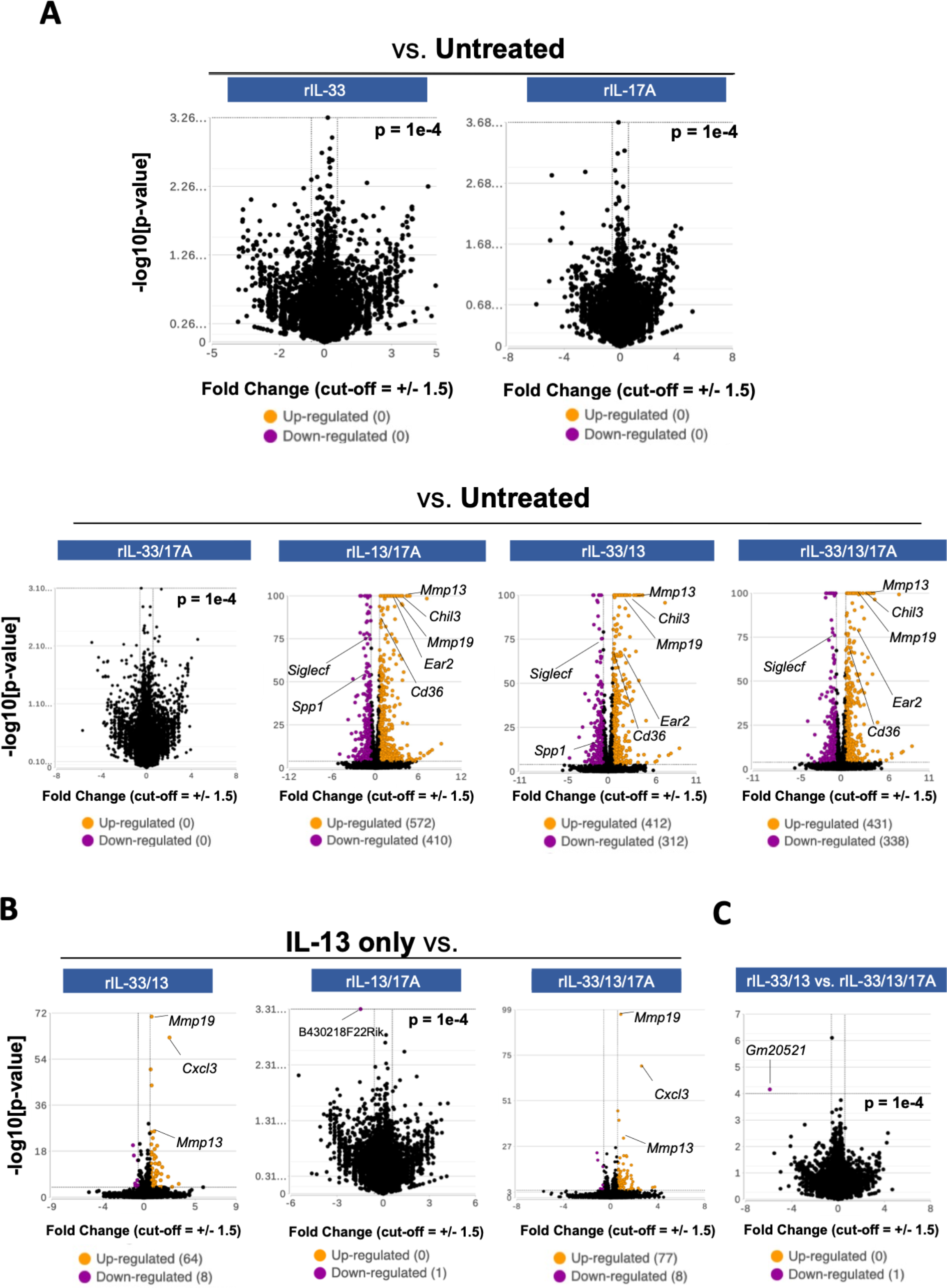
Extended transcriptional analysis of alveolar macrophages treated with combinations of recombinant IL-33, IL-13, and IL-17A relative to baseline. The algorithm DESeq2 was applied for identifying and ranking the significance of genes differentially expressed between macrophage treatment groups. DESeq2 computed an ANOVA table (fold-change, p-value, false-discovery rate step-up) for each comparison visualized here. Significantly up- or downregulated genes were illustrated by volcano plot once transformed as function of -log10[p-value] (Y-axis) and Log2[fold-change] (X-axis). Normalization of the data (median ratio) was also performed with DESeq2 by default. Significance cut-offs for this analysis was designated to be p-value <.0001, fold change = +/- 1.5. Gene identifiers in each visualization were manually annotated, and some lines are thus approximated due to size restraints and overlapping of datapoints.

**Supplemental Figure 2.**
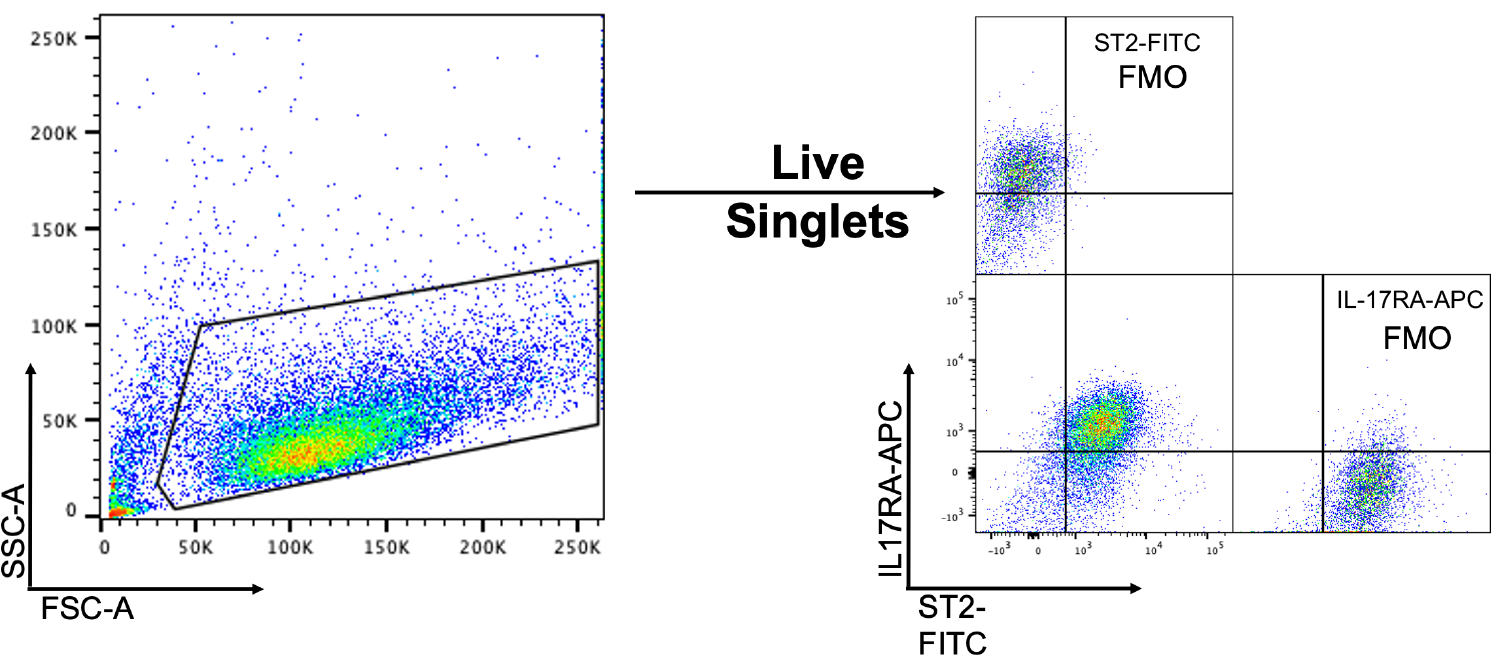
Flow cytometry for determining levels of IL17RA and ST2 on the surface of cultured lung fibroblasts. Surface protein expression measured by flow cytometry, having stained fibroblasts with anti-IL17RA-APC and anti-ST2-FITC. Positive signal for each protein was gated by comparing sample directly to a FMO control. Acquisition was performed on an LSR-II (Becton Dickinson, Franklin Lakes NJ). Cytometry of WT fibroblast was performed twice and in technical triplicate.

**Supplemental Figure 3.**
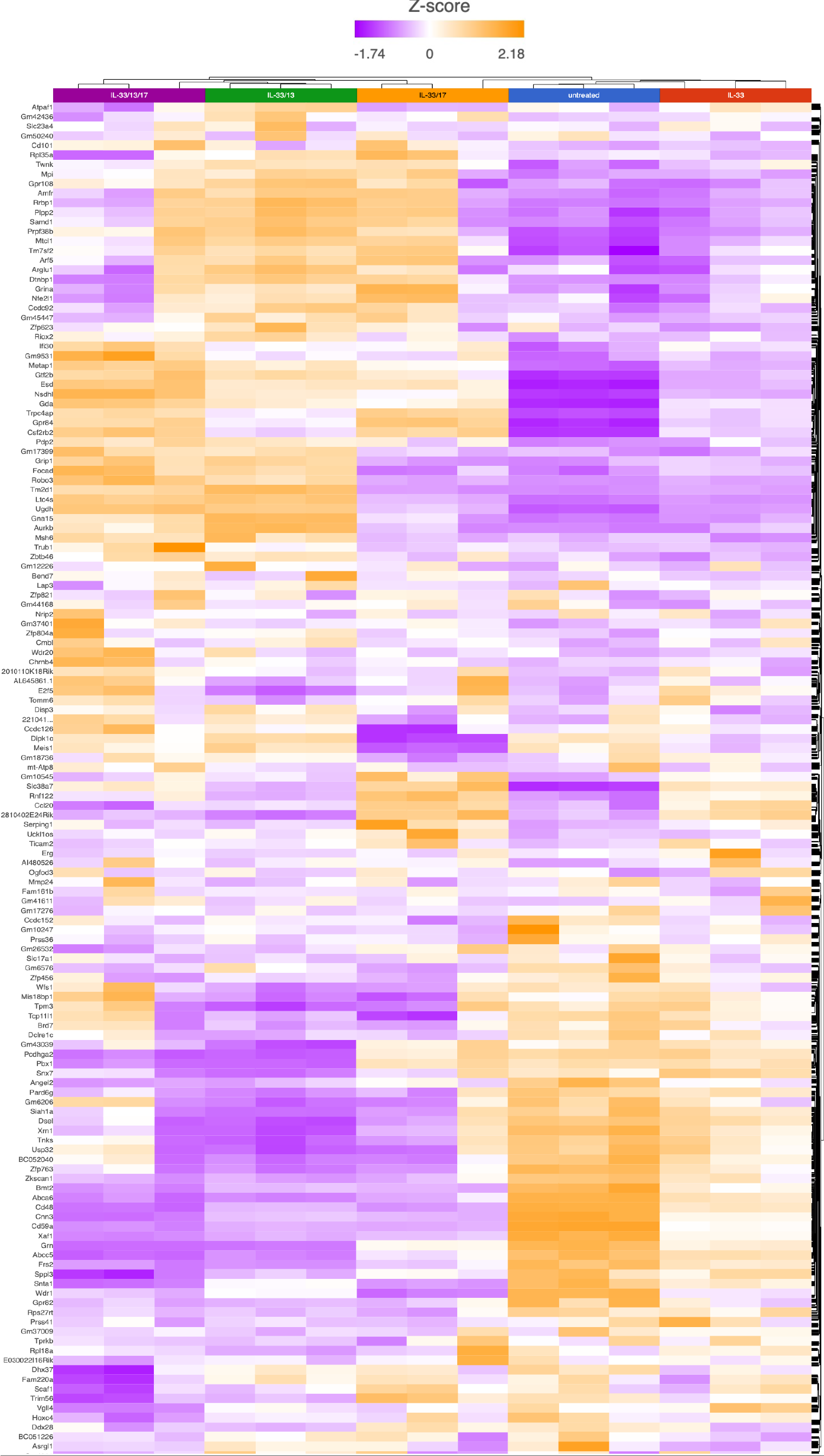
Applying hierarchical clustering for exploratory RNAseq analysis of fibroblasts stimulated with rIL-33/r-IL-13/rIL-17A. The raw count matrix first was normalized in DESeq2, which applies the Median Ratio normalization method. Treatment groups were subjected to hierarchical clustering, which projected the data as a heatmap. Columns are treatment groups and rows represent genes. Expression values for all 18,079 genes were used for computing hierarchical clustering. Though, due to size restraints, not all genes can be shown, and thus computational ‘binning’ was involved for fitting the data and generating this global representation of genes expressed per sample. Average linkage (cluster distance metric) and Pearson dissimilarity (point distance metric) were the statistical parameters applied for executing hierarchical clustering of this dataset.

**Supplemental Figure 4.**
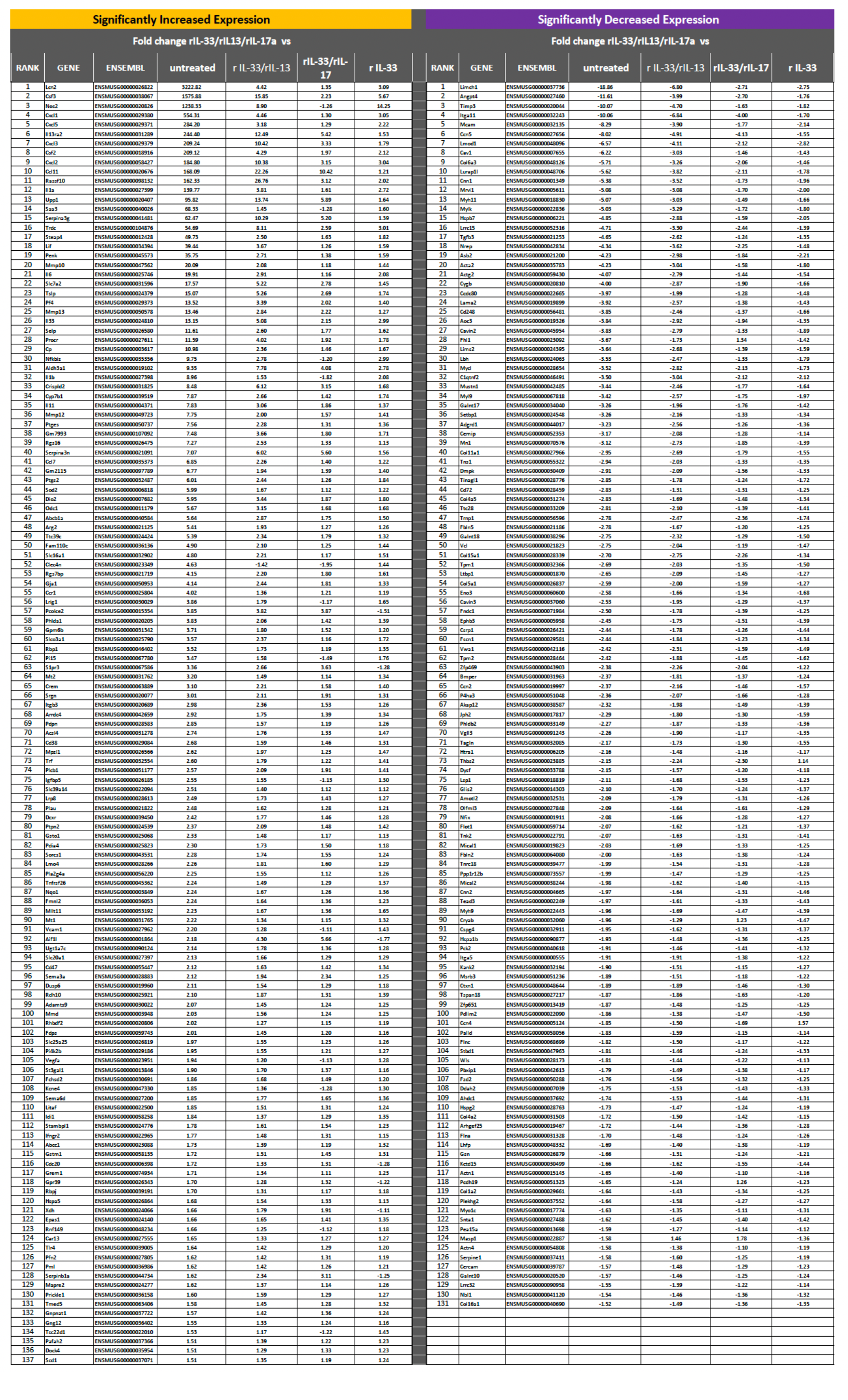
Significantly upregulated and downregulated gene expression in lung fibroblasts after rIL-33/rIL-13/rIL-17A treatment. Statistically significant (p<.0001), differential expression (fold change +/- 1.5) was noted initially for 1,642 genes (among the 18,079 detected by RNAseq) when comparing rIL- 33/13/17A-stimulated fibroblasts to untreated cells (refer to **Figure 3C**). We extended our comparative analysis by identifying statistically significant, differentially expressed genes in rIL- 33/13/17A-stimulated fibroblasts relative to every group that received cytokine treatment during the assay (*i.e.,* vs. rIL-33/rIL-13, rIL-33/rIL-17A- and rIL-33-stimulated fibroblasts). From the original list of 1,642 genes (rIL-33/rIL-13/rIL-17A-stimulated fibroblasts vs. untreated), only 268 genes were statistically significant upon comparing rIL-33/rIL-13/rIL-17A cells to each remaining treatment group (false discovery rate <.01; 131 **downregulated** and 137 upregulated). Ultimately, we ranked gene expression in this filtered ANOVA table using the **fold-change value** derived from comparing rIL-33/rIL-13/rIL-17A-stimulated fibroblasts to untreated cells.

**Supplemental Figure 5.**
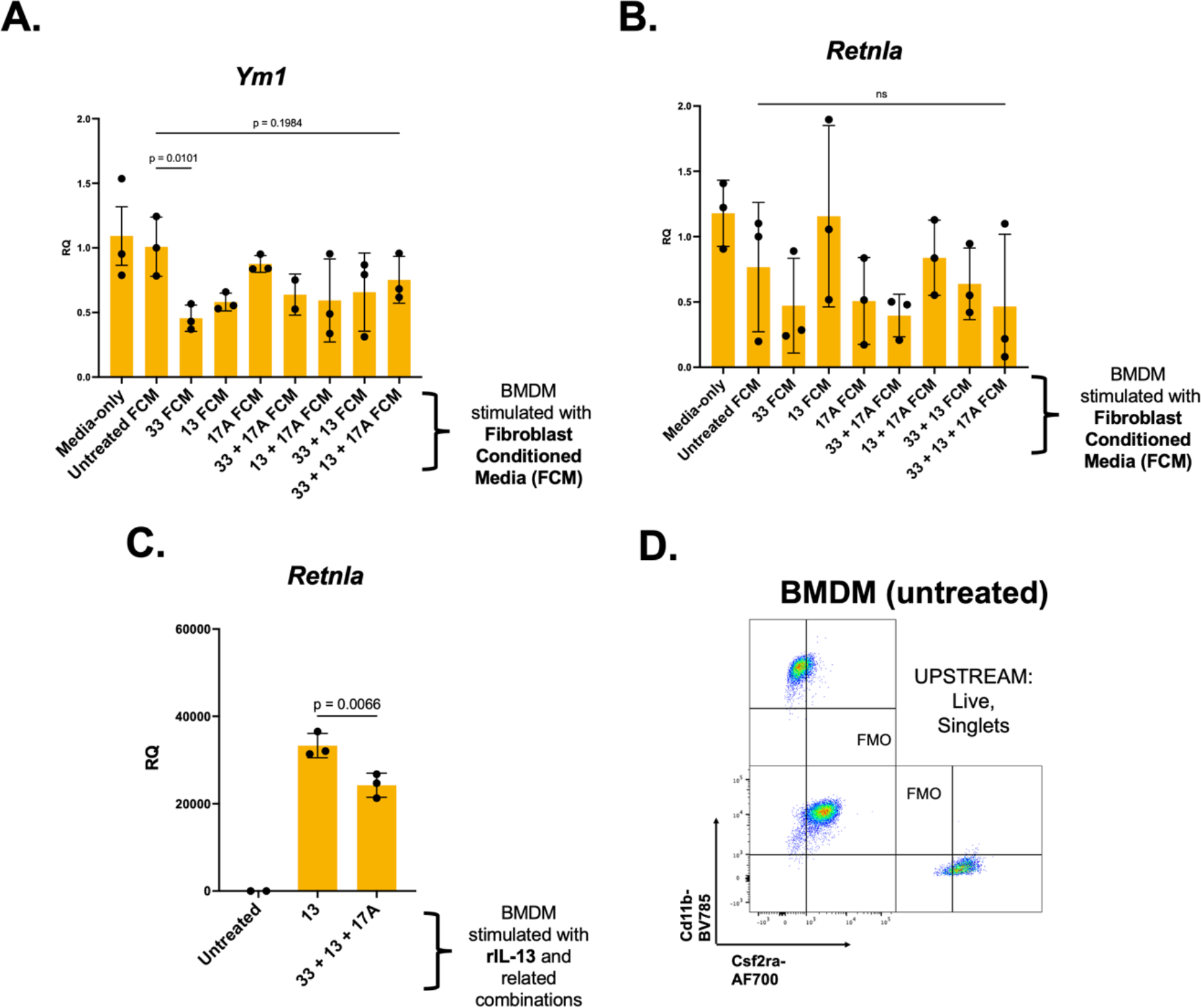
Gene and protein expression in BMDM generated for *in vitro* use. BMDM were cultured as described in **Methods**. (**A-B**) *Retnla* and *Ym1* gene expression was measured in FCM-treated BMDM at least once as part of work for supporting *Arg1* induction in these cells (refer to **Figure 5B**). There was no desire to replicate qPCR for *Ym1* and *Retnla* because nominal changes were detected upon first trial, despite normal and interpretable exponentiation of housekeeping genes assayed at the same time. (**C**) In at least one of three, conventional M2 assays performed for this work (refer to **Figure 5C**), *Retnla* was measured in BMDM treated with cytokines rIL-13 or rIL-33/rIL-33/rIL-17A directly. Significance <u>(p-value</u> < .<u>05</u>) was determined with ordinary one-way ANOVA. Multiple comparison testing was computed post-hoc, and p-values were calculated without mathematical correction using the unprotected Fischer’s Least Significant Difference test. (**A**) rIL-33/rIL-13/rIL-17A FCM vs. untreated FCM, p =0.1984; <u>rIL-33 vs. untreated FCM, p = 0.0101</u>. (**B**) ns = not significant. (**C**) <u>rIL-13 vs. rIL-33/rIL-</u> <u>13/rIL-17A, p =0.0066</u>. (**D**) Surface protein expression measured by flow cytometry, having stained BMDM with anti-CD11b-BV785 and anti-Csf2ra-Alexa Flour 700. Positive signal for each protein was gated by comparing sample directly to a FMO control. Acquisition was performed on an LSR-II (Becton Dickinson, Franklin Lakes NJ). Cytometry of WT BMDM was performed twice and in technical triplicate.

**Supplemental Table 1.**
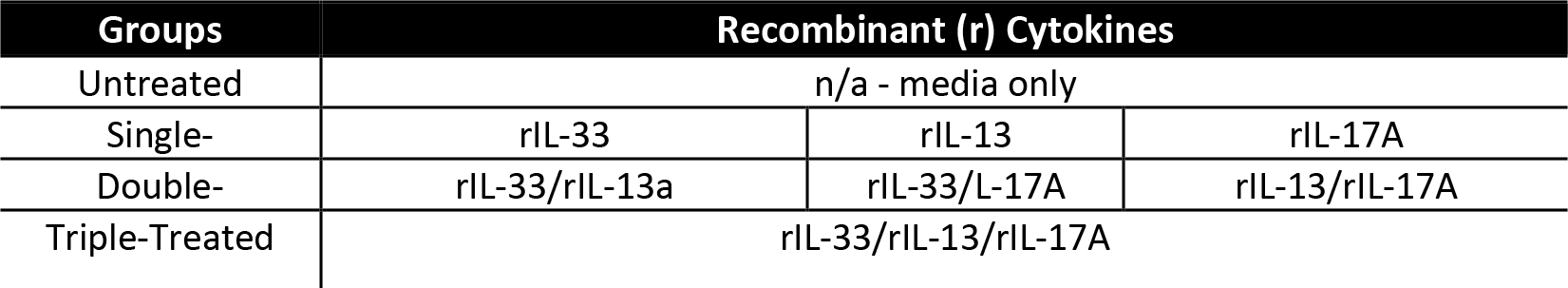
All treatment combinations for studying effects of rIL-33/rIL-13/rIL-17A *in vitro*. Recombinant cytokines were purchased from BioLegend, stored, and prepared according to manufacturer’s recommendations. Concentrations used for each study were: 30 ng/mL (rIL- 33), 20 ng/mL (rIL-13), 30 ng/mL (rIL-17A). These combinations were used for the following assays: RNAseq of alveolar macrophages and ELISA for GM-CSF secretion by primary lung fibroblasts.

| Groups | Recombinant (r) Cytokines |  |  |
| --- | --- | --- | --- |
| Untreated | n/a - media only |  |  |
| Single- | rIL-33 | rIL-13 | rIL-17A |
| Double- | rIL-33/rIL-13a | rIL-33/L-17A | rIL-13/rIL-17A |
| Triple-Treated | rIL-33/rIL-13/rIL-17A |  |  |

**Supplemental Table 2.** Cytokine combinations used for RNAseq of primary lung fibroblasts. Recombinant cytokines were purchased from BioLegend, stored, and prepared according to manufacturer’s recommendations. Concentrations used for this study were: 30 ng/mL (rIL- 33), 20 ng/mL (rIL-13), 30 ng/mL (rIL-17A).

| Groups | Recombinant (r) Cytokines |  |
| --- | --- | --- |
| Untreated | n-a - media only |  |
| Single- | rIL-33 |  |
| Double- | rIL-33/rIL-13 | rIL-33/L-17A |
| Triple-treated | rIL-33/rIL-13/rIL-17A |  |

